# Genome-wide association study of piscine myocarditis virus (PMCV) robustness in Atlantic salmon (Salmo salar)

**DOI:** 10.1101/458901

**Authors:** Borghild Hillestad, Hooman K. Moghadam

## Abstract

Cardiomyopathy syndrome is a sever, viral disease of Atlantic salmon that mostly affects farmed animals during their late production stage at sea. Caused by piscine myocarditis virus (PMCV), over the past few years, the outbreaks due to this disease have resulted in significant losses to the aquaculture industry. However, there are currently no vaccine that has proven effective against this virus. In this study, using a challenge model, we investigate the genetic variation for robustness to PMCV, by screening large number of animals using a 55 K SNP array. In particular, we aimed to identify genetic markers that are tightly linked to higher disease resistance and can potentially be used in breeding programs. Using genomic information, we estimated heritability of 0.41 ±0.05, suggesting that robustness against this virus is largely controlled by genetic factors. Through association analysis, we identified a major QTL on chromosome 27, explaining approximately 57% of the total additive genetic variation. The region harbouring this putative QTL contains various immune related candidate genes, many of which have previously been shown to have a differential expression profile between the naïve and infected animals. We also identified a suggestive association on chromosome 12, where the QTL linked markers are located within two putatively immune related genes. These findings are important as they can be readily implemented into the breeding programs but also the results can further help in fine-mapping the causative mutation, in better understanding the biology of the disease and refine the mechanics of resistance against PMCV.

## Introduction

Cardiomyopathy syndrome (CMS) is an emerging, severe inflammatory cardiac disease of Atlantic salmon, with the farmed animals being the primary host, although the causative agent has also been detected in wild populations (Brun *et al.*, 2003; Garseth *et al.*, 2012). The disease has been linked to piscine myocarditis virus (PMCV), a double stranded RNA virus that resembles members of the *Totiviridae* family (Løvoll *et al.*, 2010; Haugland *et al.*, 2011). First detected in farmed Atlantic salmon in Norway in 1985 (Amin and Trasti, 1988), outbreaks have since been reported in other geographical regions and countries including the Faroe Islands, Scotland, Ireland and possibly Canada (Rodger and Turnbull, 2000; Brocklebank and Raverty, 2002; Garseth *et al.*, 2018). This disease is of major concern since it can pose significant financial burden to both farmers and to the industry, as it usually affects the brood fish or grow-out fish at their later life stages in sea (Svendsen and Fritsvold, 2009; Garseth *et al.*, 2018), where considerable amount of recourses have already been allocated for the rearing and husbandry of the animals. It seems however, that the frequency of younger fish also being diagnosed by this virus might be on the rise (Svendsen and Fritsvold, 2018). An initial assessment in 2003, indicated that the cost of CMS associated outbreaks are responsible for direct annual financial loss of € 4.5 to 8.8 million to the Norwegian aquaculture sector (Brun *et al.*, 2003). However, a subsequent study in 2011, published by The Norwegian Seafood Research Fund (FHF), estimated that the overall CMS related losses in 2007 approximated to more than € 25 million (FHF, 2011). These estimates suggest that the financial loss due to this disease has increased by four- to five-fold in mere four years. According to a recent report by the Norwegian Veterinary Institute (Svendsen and Fritsvold, 2018), the frequency of CMS outbreaks has increased throughout Norway over the past few years. Currently, CMS is considered as one of the most significant health related challenges to the aquaculture industry, ranking only after the salmon-lice and gill-diseases (Svendsen and Fritsvold, 2018).

A few studies have so far investigated the virus at its genomic details. PMCV has a small genome of 6,688 nucleotides, consisting of three open reading frames (ORF) (Løvoll *et al.,* 2010; Haugland *et al.*, 2011). While the putative products of the ORF1 and ORF2 are likely to be involved in encoding the protein coat and RNA-dependent RNA polymerase respectively, the exact functional properties of ORF3 product is yet to be determined (Wiik-Nielsen *et al.*, 2013). Analysis of the Norwegian PMCV isolates, collected from 36 farms, has revealed high similarity in the nucleotide sequence information, with the most divergent isolate sharing more than 98% sequence similarity (Wiik-Nielsen *et al.*, 2013). Sequence analysis of a few Irish (Rodger *et al.*, 2014) as well as wild Atlantic salmon isolates from Norway (Garseth *et al.,* 2012) have also shown very high similarity to the Norwegian variants. This suggest that all these subtypes, most likely belong to a single genus. Although our understanding of the dynamics and biology of CMS is still very limited, a few studies have started to shed light on the host transcriptomic response following infection with PMCV. In particular, Timmerhaus *et al.* (2011), through comparative analysis of gene expression data and assessment of histopathological lesions in different timepoints and tissues, have identified alternative regulation of six different gene sets during the course of infection. These gene sets included i. genes involved in early antiviral and interferon response; ii. complement response; iii. B cell response; *iv.* MHC antigen presentation; v. T cell response and *vi.* apoptosis. Further, studies have reported large, inter-individual variation in Atlantic salmon’s response to PMCV, regarding both the progression and the pathological outcomes of the disease (Timmerhaus *et al.*, 2012; Garseth *et al.*, 2018). While some fish seemed to be able to clear or significantly reduce the level of the virus from 6-10 weeks post infection and exhibit little evidence of disease pathology, other fish retained high loads of virus and elevated heart tissue damage (Timmerhaus *et al.*, 2012). The authors also noticed that at the final stages of the challenge, a broad range of immune related genes, genes, mainly involved in adaptive immunity and particularly in T cell response, had altered their profile of expression among more susceptible animals (Timmerhaus *et al.*, 2012). Therefore, one might speculate that a significant part of the resistant machinery against this virus should be under genetic control (Garseth *et al.,* 2018), with genes involved in adaptive immunity to most likely play an important role in this process.

In the absence of any effective vaccine or available treatment for CMS (Garseth *et al.*, 2018), alternative strategies such as selective breeding and utilization of the latest genomic technologies and genetic recourses can provide us with innovative ways to help in identifying resistant or more tolerant animals, reduce the frequency of the disease outbreak, help to improve animal welfare and increase the profit to the industry. With such goals in mind, in this study, we aimed to identify chromosomal regions and genomic markers that are associated with resistance or higher tolerance against PMCV. We further investigated the heritability of the trait and suggest a number of candidate genes that might harbour the causative variation(s) and might be responsible for making an animal to cope better with the pathological symptoms of this virus.

## Materials and Methods

### SalmoBreed population and challenge test

In the Fall of 2017, 1,192 PIT-tagged (passive integrated transponder) smolts were transported from the SalmoBreed breeding station in Lønningdal (http://salmobreed.no/en/) to the challenge facility in VESO Vikan (https://www.veso.no/about-us1; Namsos, Norway). The group consisted of 60 full-sib families, approximately 20 individuals per family, from the SalmoBreed nucleus year-class 2017, with an average weight of 123 gr. After arrival, 1,186 fish were kept at 12 °C brackish water (15-30%o) and 24:00 h light regime. After the initial acclimatization, the remaining 1,179 fish were challenged with PMCV in a full salinity water (>30‰). The fish were first anaesthetized, scanned and then challenged through intraperitoneal injection (0.1 mL per fish), with a virus containing tissue homogenate, cultured *in vivo* at VESO Vikan. The challenge was carried out for nine weeks before the trial was terminated. At termination, the fish were weighted, and heart tissue samples were collected for histopathology and quantitative real-time PCR (qRT-PCR) analyses, and stored in formalin and RNALater, respectively. The adipose fin tissue was also collected for DNA extraction and subsequent genotyping.

### RNA extraction and qRT-PCR analysis

Following the termination and collection of the heart tissues in RNALater, samples were shipped to PatoGen AS (http://www.patogen.com/; Ålesund Norway) for RNA extraction and viral load quantification using their established protocol. RNA was successfully extracted from 1,161 heart tissue samples. Real-time qRT-PCR of the viral loads and calculations of normalized cycle threshold (Ct) values were performed based on the optimized procedure for PMCV quantification in PatoGen.

### Histology assessment

To estimate the correlation between *Ct* values of the viral loads from qRT-PCR and heart histopathology, 40 fish where selected for histology assessment of both atrium and ventricle at the Fish Vet Group Norway (http://fishvetgroup.no/en/; Skøyen, Norway). Formalin-fixed heart tissue samples were embedded in paraffin and processed in accordance to the Fish Vet Group’s routine standard histological procedures (Bott, 2014). Histology analysis were performed on 20 fish with the lowest and 20 fish with the highest *Ct* values. The lesions were scored 0, 1, 2 or 3 in accordance with the scheme described previously (Timmerhaus *et al.*, 2011). A score of 0 indicates no lesions at the heart, 1 refers to mild lesions, 2 states moderate lesions and 3 implies severe lesions.

### Genotyping and genotype quality assessment

The adipose fin-clip tissue from all the animals that survived the nine weeks duration of the challenge (i.e., 1,182 individuals) were sent to IdentiGEN (https://identigen.com/; Dublin, Ireland) for DNA extraction and genotyping. The genotyping was done on a custom made 55 K Affymetrix Axiom array, called N0FSAL03, developed by Nofima AS in 2016 in collaboration with SalmoBreed AS and Marine Harvest ASA. In total, 1,152 fish passed the initial quality control during DNA extraction and the SNP calling steps of the Affymetrix Axiom analysis suite software. Additional genotype quality measures were undertaken using SNP & Variation Suite v8.8.1 (SVS; Golden Helix Inc., Bozeman, MT, USA www.goldenhelix.com). Samples and SNPs with call rates < 90%, SNPs with Hardy-Weinberg p-value (Fishers exact test) < 10^-10^ or genetic markers with minor allele frequency < 0.05% were excluded from downstream analysis. Further, we used SVS for linkage-disequilibrium pruning, by setting window size to 40, window increment to 5 and *r^2^* threshold to 0.5. We then used the pruned data to construct a distance matrix based on the identity by descent (IBD). Using this pre-computed kinship matrix, we performed mixed model association analysis by applying Efficient Mixed-Model Association eXpedited (EMMAX) (Kang *et al.*, 2010) to correct for possible sample structures and relatedness between animals. An association was considered to be genome-wide or chromosome-wide significant, if the Bonferroni threshold *p*-value was less or equal than 1.004e-06 or 3.730e-05 respectively.

To estimate the proportion of total genetic variation that is explained by genomic loci harboring QTL, we performed regional heritability analysis (Nagamine *et al.,* 2012) using DISSECT (Canela-Xandri *et al.*, 2015). We first extracted the SNPs covering the QTL regions of interest and computed a genetic relationship matrix between individuals *i* and *j* as:

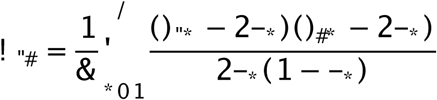

where *S_ik_* and *S_jk_* are the number of copies of the reference allele for SNP *k* in individuals *i* and j, *p_k_* is the frequency of the reference allele for SNP *k* and *N* is the number of SNPs. We then fitted a mixed linear model as follows:

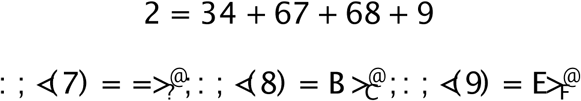

where *y* is the vector of *Ct* values, *X* and *Z* are the design matrixes for fixed and random effects respectively, *u* is the global genomic additive genetic effect, *v* is the regional genomic additive genetic effect, *e* is the residuals and *β* is the fixed effect, adjusted for sex. Matrices *G* and *I* are a global genomic relationship matrix using the entire quality control passed SNPs for estimating global genomic additive effect and a unit matrix for estimating residuals, respectively. *Q* is the regional genomic relationship matrix obtained from the SNPs covering the QTL region. The global and the regional heritability are estimated as 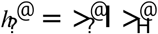 and 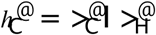 respectively.

## Results and Discussion

Following the termination of the trial, no mortality or clinical symptoms associated with the disease was observed. Using qRT-PCR, the load of the virus was successfully obtained from the heart tissues of 1,169 animals. The standardized *Ct* values of the viral genetic material showed a normal distribution when compared across individuals, with some animals exhibiting very low levels of the pathogen while some others carrying high loads of PMCV (Supplementary Figure 1). This is an indication of possible inter-individual biological differences in the ability of an animal to either prevent the proliferation or subsequent clearance of the virus during later stages of infection (Timmerhaus *et al.*, 2011, 2012). To test if these quantitative measures can be used as a proxy for assessing the degree of the damage to the heart and therefore providing an indication of an animal’s resistance or tolerance against PMCV, histopathological assessments were performed on the atriums and ventricles of 40 animals, 20 with the highest and 20 with the lowest *Ct* measurements. The heart histopathological scores ranged from 0-2 (no score of 3 was observed), reflecting the degree of severity of the lesions as described previously (Timmerhaus *et al.*, 2011). The *Ct* values were highly correlated with the histology scores, for both atrium and ventricle (Pearson *r* = 0.72,*p*-value < 0.0001; Figure 1). Similar, high estimates of correlation (i.e., 0.75-0.76) between the histopathological scores and the loads of virus have also been previously reported for this disease (Haugland *et al.*, 2011; Timmerhaus *et al.*, 2011).

**Figure 1.**
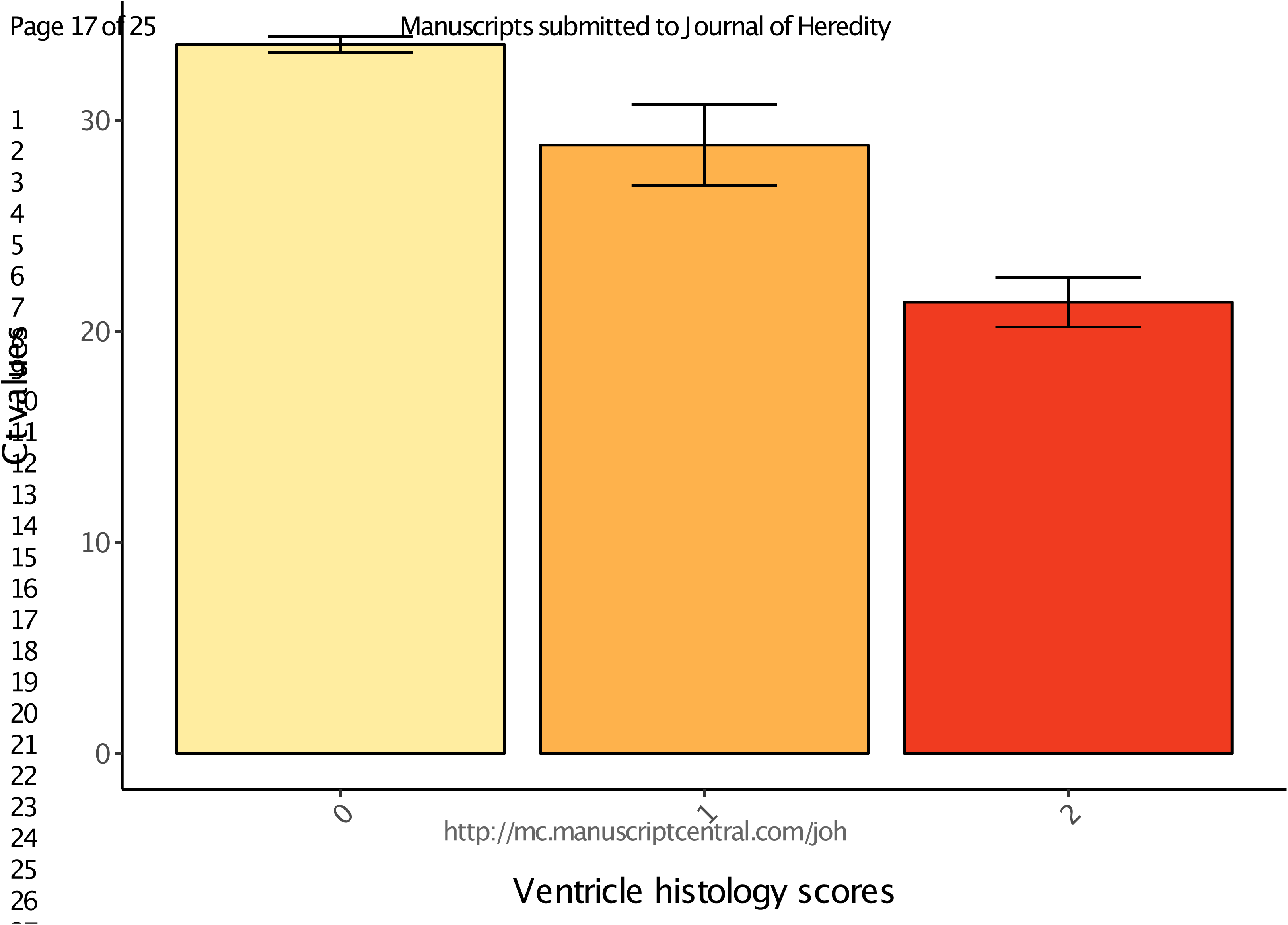
qRT-PCR *Ct* values and the associated standard errors of the PMCV viral-loads in animals with ventricle histology scores of 0, 1 and 2.

To identify genetic markers that are associated to robustness against PMCV, we performed a genome-wide association study (GWAS). The markers were first filtered for minor allele frequencies < 0.05 and call-rates < 90%, leaving approximately 50 K SNPs for the subsequent analysis. The test was preformed using EMMAX algorithm, in order to account for both animal relatedness as well as population structure (Kang *et al.*, 2008). We obtained an inflation factor (/) of 1.13, suggesting a negligible population structuring effect in our data (Supplementary Figure 2). The heritability estimate, using the genomic relationship matrix, was 0.41 ± 0.05, indicating potentials for efficient response in increased resistance against this pathogen through selection and breeding strategies. Through GWAS analysis, we identified a significant association on chromosomes 27 (-log_10_*p*-value > 6) but also a suggestive evidence of genetic markers on chromosome 12 to be linked with higher robustness against PMCV (-log_10_ *p*-value > 4.5) (Figure 2). On chromosome 27, total of 44 SNPs passed the genome-wide threshold *p*-value of 1.004e-06 (Supplementary Table 1). These markers cover a large fragment on the chromosome, spanning from 3.9 Mbp up-to 25.5 Mbp region. However, the top four SNPs, with a much stronger association compared to the remainder of the genetic markers, cover only 2 Mbp segment of the chromosome, from 8.5 Mbp to 10.5 Mbp. The proportion of phenotypic variance explained by each of these four markers were 5.13%, 5.03%, 4.81% and 4.05% respectively. Collectively, the markers within this 2 Mbp region explained about 57% of the total additive genetic variation, according to the regional heritability estimate.

**Figure 2.**
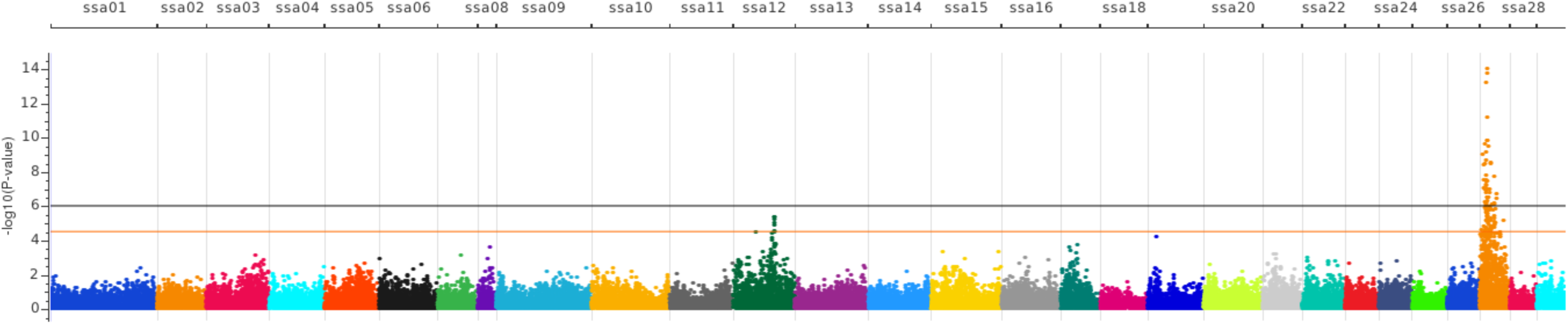
Manhattan plot of association between genetic markers and cardiac viral-loads of PMCV in Atlantic salmon. The black and the orange lines indicate the genome-wide and the chromosome-wide significance threshold cut-off levels respectively.

This region of chromosome 27 in Atlantic salmon harbors about 63 protein coding genes, with the strongest associated SNP located on 10.4 Mbp (ssa27:10393267; Table 1). On the downstream of this SNP, there are two putative immune related candidate genes, immunoglobulin V-set-like domain (L0C106588384; ssa27:10,413,985-10,428,204) and the major histocompatibility complex class I-related gene (L0C106588381; ssa27:10,464,840-10,482,970). These two genes have potential functions in antigen binding and antigen presentation respectively. On the upstream of this SNP, there are further a few genes with direct functional properties relevant to the immune system processes and antigen presentation. Antigen peptide transporter 2 (*tap2b*; ssa27:10,176,035-10,180,872), class I histocompatibility antigen F10 alpha chain-like (L0C106588401; ssa27:10,122,008-10,149,393), major histocompatibility complex class I-related gene (L0C106588402; ssa27:10,037,565-10,054,429), TAP-binding protein (*tapbp*; ssa27:10,023,358-10,039,557) and proteasome subunit beta type-7-like (L0C106588382; ssa27:10,580,776-10,585,552) are a few examples. All these latter genes cover a region on chromosome 27 from 10 Mbp up-to 10.2 Mbp. Previously, Timmerhaus *et al.* (2011), using a microarray platform, have identified a set of 34 transcripts that are involved in the presentation of viral antigens through MHC class I and II genes and were differentially expressed between the naïve and the CMS infected animals. In fact, we found that the genes associated with a few of these transcripts are located within the CMS QTL genomic segment on chromosome 27 reported in this study. In particular, these genes included L0C106588401, L0C106588402 (as indicated above), and to a lesser degree characterized gene, L0C106588388 (ssa27:10,583,372-10,595,992), with possible functionality as a long non-coding RNA (lncRNA).

Further analysis of the 34 transcript set of the sequence data reported by Timmerhaus *et al.* (2011) also showed that a subset of these transcripts further map to both chromosomes 14 and 27 in the current Atlantic salmon genome assembly (GCA_000233375.4 ICSASG_v2). Examples of these transcripts mapping to the two chromosomes include proteasome subunit beta and MHC class I. In Atlantic salmon, the entire chromosome 27 is homeologous to chromosome 14, suggesting that these chromosomes have originated from the whole genome duplication event that happened at the origin of all salmonid fishes (Lien *et al.*, 2016). On chromosome 27, the duplicates reported by Timmerhaus *et al.* (2011) mainly fall within the QTL associated genomic segment. While the exact functional properties of many duplicated gene copies in Salmonids have yet to be studied in detail, we expect that many genes to have either developed new functions (*neojunctionalization*) (Berthelot *et al.*, 2014) or the ancestral function has been partitioned between the two newly derived daughter copies (*subjunctionalization*) (Wolfe, 2001; Osborn *et al.*, 2003). In addition, it is possible that both copies have remained equally functional within the genome, if the dosage effect provides an advantage and increases the fitness of an individual (Wolfe, 2001). Therefore, it is of interest to find out if in some other Atlantic salmon populations, the homeologous segment on chromosome 14, either by itself or in addition to chromosome 27, to show association with higher PMCV robustness. Further, in the future studies, targeting to investigate the expression and the exact genomic location of these differentially expressed transcripts, whether it is on chromosome 14 or 27, between animals with different levels of resistance to CMS, would be of great importance, particularly when we aim to fine-map the causal mutation for this trait.

On chromosome 12, while no marker passed the genome-wide, threshold *p*-value, eight markers exceeded the chromosome-wide significance corrected threshold of 3.730e-05 (Supplementary Table 1). Seven of these markers span a region of only 320 Kbp, from position 61.39 Mbp to 61.71 Mbp. The proportion of phenotypic variance explained by these markers range from 1.48% to 1.83% (Supplementary Table 1). There are 15 annotated, protein-coding genes within 61.00 to 62.00 Mbp segment of chromosome 12. However, probably the two most relevant and plausible candidate genes, within this region, with a potential effect on an animal’s robustness to PMCV are the two putative H-2 class II histocompatibility antigen genes. One of these genes, L0C100136577, is homologous to H-2 class II histocompatibility antigen, I-E beta chain in mouse, located at ssa12:61,693,946-61,699,456. The other gene, L0C106565699 is most similar to H-2 class II histocompatibility antigen, A-U alpha chain, again in mouse, located at ssa12:61,701,374-61,703,966. In fact, four of the SNPs with the lowest associated *p*-values detected on this chromosome, are located within this latter gene. Two of the SNPs fall within the intronic regions of the gene, while the other two have been assigned as 3Cuntranslated region (UTR) variants (Figure 3). Interestingly, this genes has also been reported by Timmerhaus *et al.* (2011) as one of the key genes, within the MHC antigen presentation set, that is differentially expressed between the PMCV infected and non-infected animals. The 5^th^ strongest SNP on chromosome 12, falls in the other histocompatibility gene (i.e., L0C100136577) and has also been identified as a 3¢CUTR variant (Figure 3). These five SNPs are in strong linkage disequilibrium and form a haplotype block.

**Figure 3.**
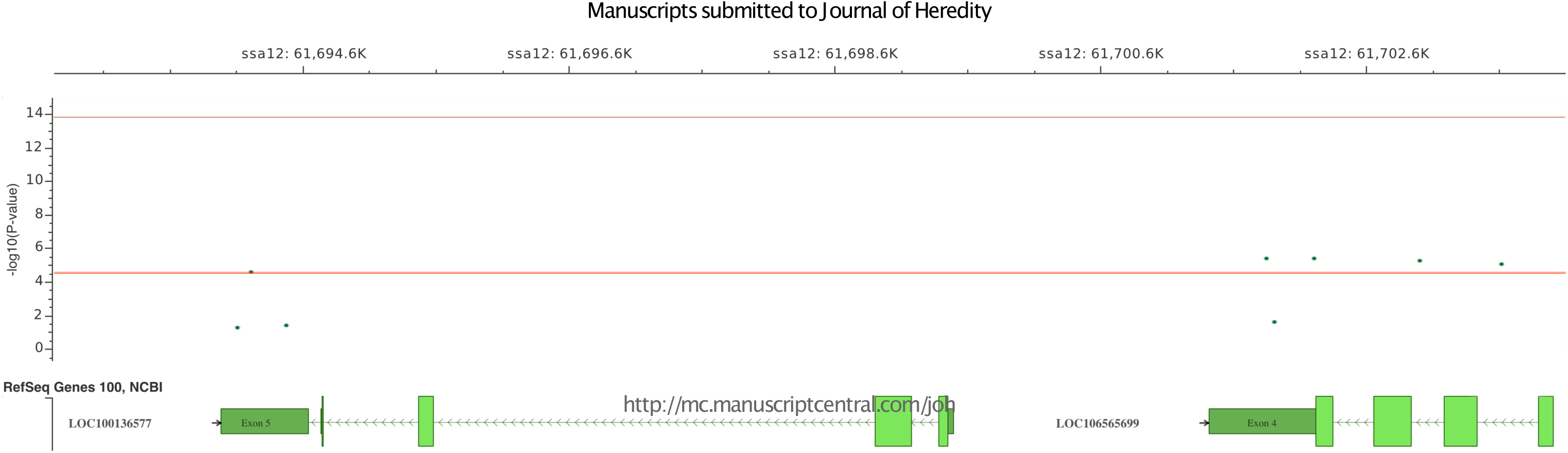
Chromosome-wide significant genetic markers on chromosome 12, located on two putative H-2 class II histocompatibility antigen genes, L0C100136577 and L0C106565699. The darker green blocks represent the UTR regions of the gene while the light green blocks represent the protein coding sequences. The arrowed lines show the intronic segments of the gene.

In addition to the two histocompatibility genes however, there are also other putative candidate genes with direct or indirect functional properties relevant to the immune system, within this approximately 320 Kbp segment of the chromosome. One example is the T cell transcription factor EB-like gene (TFEB) (LOC106565694), which can be regarded as one of the key transcriptional regulators of autophagy and lysosome biogenesis (Sardiello *et al.*, 2009; Settembre *et al.*, 2011). This gene is located in close proximity to the two histocompatibility genes discussed above (ssa12:61,588,376-61,651,838). The product of this gene is involved in a variety of functions which are related to the host’s defense mechanism, including elimination of intracellular pathogens, reducing inflammation, antigen presentation and secretion of cytokines (Nabar and Kehrl, 2017).

The final, chromosome-wide significant SNP detected on chromosome 12, is located further apart from the other associated markers, at approximately 33 Mbp location (Supplementary Table 1). Interestingly, this marker is within a putative long-noncoding RNA (lncRNA, LOC106565045, ssa12:33,241,465-33,247,235). Long-noncoding RNAs are known to be involved in gene transcription, translation and regulation and their key roles in many disease progressions have previously been suggested (Carrieri *et al.*, 2012).

In conclusion, this work provides an important first step towards unraveling the genetic architecture of resistance against PMCV. Identification of a major QTL on chromosome 27 that explains a large proportion of genetic variation and suggestive evidence of one or more genomic regions on chromosome 12 that are associated with CMS robustness indicates that breeding can be a powerful tool for reducing and managing outbreaks due to this virus. It is important however, to confirm the same association of the QTL in different year-classes and in different populations. Further, understanding the genes and the genetic networks that are differentially regulated between the resistant and susceptible animals, particularly the expression profile of those genes that fall within the QTL region will help in fine mapping and identifying the causative mutation. As obtaining the whole-genome sequence information from an increasing number of animals is becoming more affordable, this will be a key and a routine step in our future attempts in dissecting the genetic basis of any complex trait.

## Acknowledgments

The authors would like to thank Dr. Makoto Inami and other personnel at the VESO Vikan who helped in the development and optimization of the CMS challenge test. We would also like to extend our gratitude to the Fish Vet Group, Norway and specially Dr. Kai-Inge Lie for performing histology on the heart tissue samples.

---

**Supplementary Figure 1.**
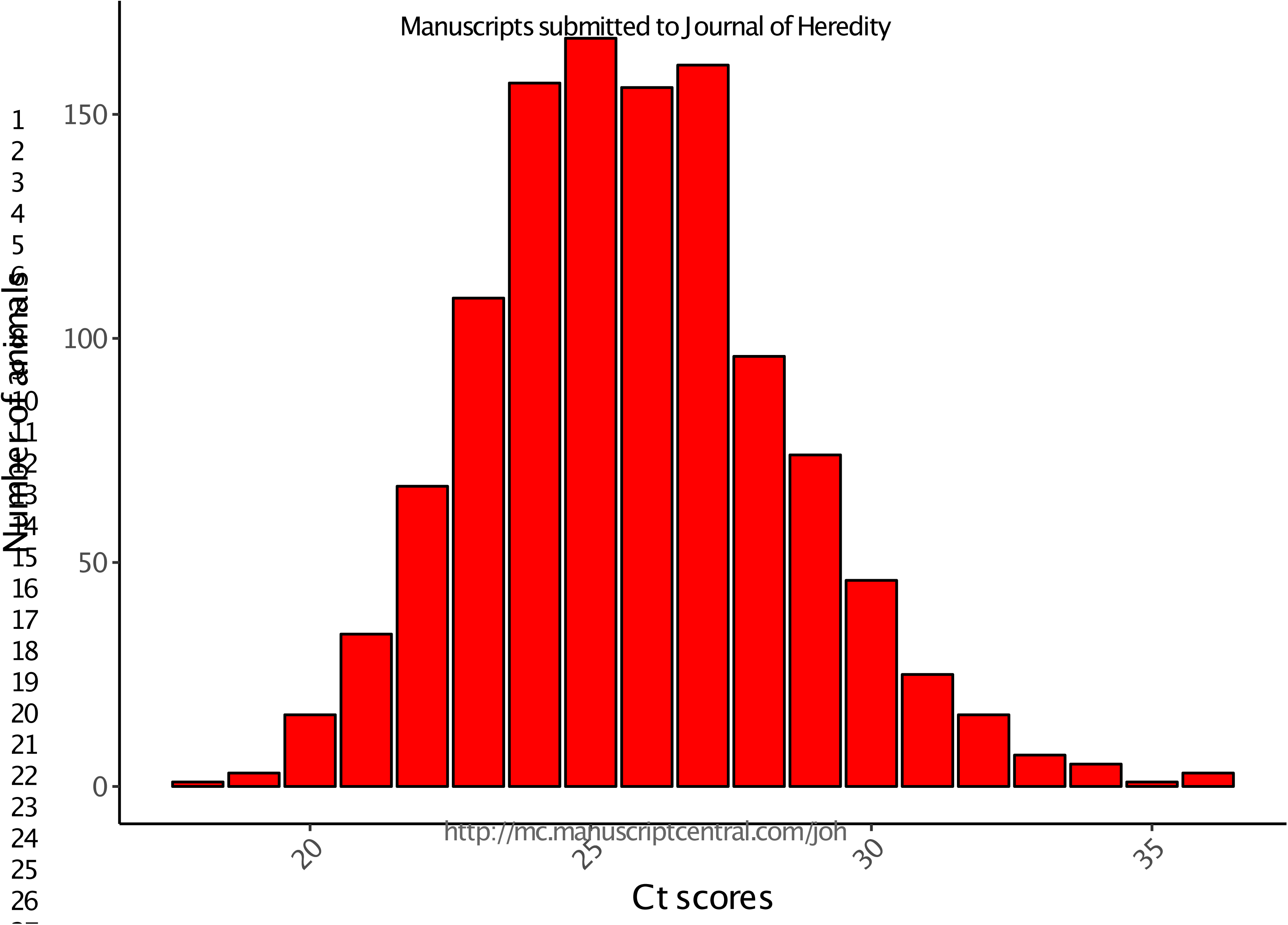
Bar-graph showing the distribution of the normalized viral-loads *Ct* values from qRT-PCR across all challenged animals.

**Supplementary Figure 2.**
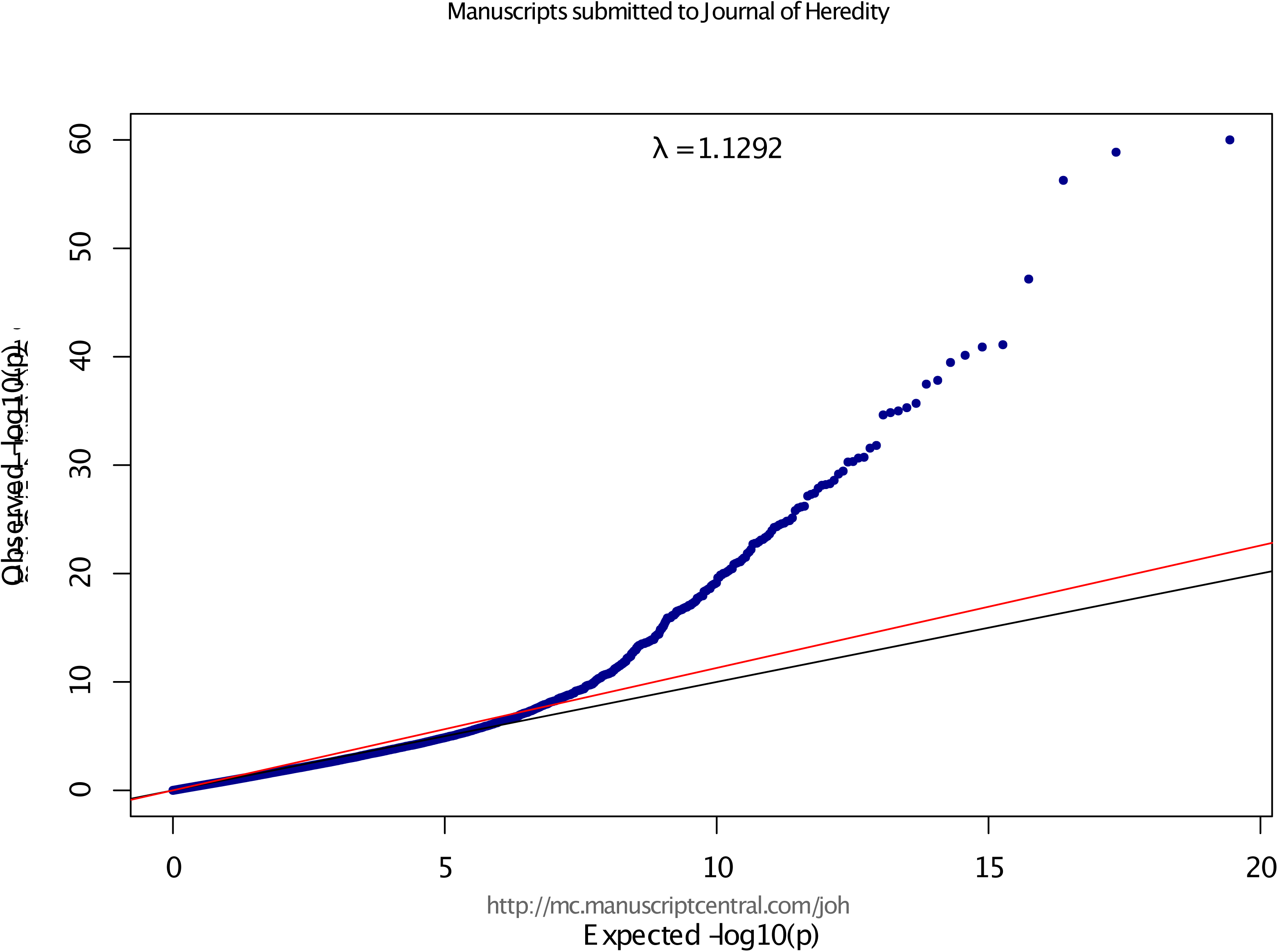
Quantile-quantile plot for the genome-wide association analysis for robustness to PMCV.

**Supplementary Table 1.** List of genome-wide and chromosome-wide genetic markers associated with robustness against PMCV in Atlantic salmon.

| Marker | Chromosome | Position | *p*-value | $-\log_{10} p$-value | Proportion of variance explained | Significance |
| --- | --- | --- | --- | --- | --- | --- |
| AX-87431771 | ssa27 | 2703017 | 2.32E-05 | 4.63 | 1.56% | Chromosome-wide |
| AX-88206216 | ssa27 | 3921804 | 9.27E-10 | 9.03 | 3.23% | Genome-wide |
| AX-98319434 | ssa27 | 3992934 | 3.62E-05 | 4.44 | 1.48% | Chromosome-wide |
| AX-88070502 | ssa27 | 4706791 | 3.98E-09 | 8.40 | 2.99% | Genome-wide |
| AX-88113080 | ssa27 | 5028120 | 8.89E-08 | 7.05 | 2.48% | Genome-wide |
| AX-87777813 | ssa27 | 5747851 | 1.61E-05 | 4.79 | 1.62% | Chromosome-wide |
| AX-88060928 | ssa27 | 6292856 | 3.09E-08 | 7.51 | 2.65% | Genome-wide |
| AX-96121015 | ssa27 | 6912223 | 1.05E-07 | 6.98 | 2.45% | Genome-wide |
| AX-87957864 | ssa27 | 7349971 | 1.29E-06 | 5.89 | 2.03% | Chromosome-wide |
| AX-87770393 | ssa27 | 7558502 | 3.57E-09 | 8.45 | 3.01% | Genome-wide |
| AX-87810183 | ssa27 | 7595841 | 2.37E-10 | 9.63 | 3.46% | Genome-wide |
| AX-88146773 | ssa27 | 7778033 | 2.27E-05 | 4.64 | 1.56% | Chromosome-wide |
| AX-87543614 | ssa27 | 7891594 | 6.09E-07 | 6.22 | 2.16% | Genome-wide |
| AX-88290504 | ssa27 | 8039250 | 3.03E-05 | 4.52 | 1.51% | Chromosome-wide |
| AX-88223822 | ssa27 | 8282588 | 9.85E-07 | 6.01 | 2.08% | Genome-wide |
| AX-87347259 | ssa27 | 8356623 | 2.30E-09 | 8.64 | 3.08% | Genome-wide |
| AX-87353154 | ssa27 | 8372617 | 1.68E-06 | 5.77 | 1.99% | Chromosome-wide |
| AX-98308488 | ssa27 | 8374637 | 1.30E-07 | 6.89 | 2.41% | Genome-wide |
| AX-88285513 | ssa27 | 8617575 | 2.97E-08 | 7.53 | 2.66% | Genome-wide |
| AX-88190840 | ssa27 | 8642177 | 6.30E-14 | 13.20 | 4.81% | Genome-wide |
| AX-87513090 | ssa27 | 8972988 | 6.61E-08 | 7.18 | 2.53% | Genome-wide |
| AX-88286284 | ssa27 | 8979164 | 1.69E-08 | 7.77 | 2.75% | Genome-wide |
| AX-105314687 | ssa27 | 9008115 | 3.73E-08 | 7.43 | 2.62% | Genome-wide |
| AX-87791679 | ssa27 | 9056093 | 6.26E-07 | 6.20 | 2.15% | Genome-wide |
| AX-98308480 | ssa27 | 9056598 | 7.52E-07 | 6.12 | 2.12% | Genome-wide |
| AX-87067374 | ssa27 | 9056780 | 6.63E-06 | 5.18 | 1.76% | Chromosome-wide |
| AX-87003873 | ssa27 | 9233208 | 5.74E-08 | 7.24 | 2.55% | Genome-wide |
| AX-87517328 | ssa27 | 9287317 | 1.74E-07 | 6.76 | 2.37% | Genome-wide |
| AX-88202021 | ssa27 | 9297577 | 2.43E-06 | 5.61 | 1.93% | Chromosome-wide |
| AX-98321803 | ssa27 | 9308180 | 7.74E-10 | 9.11 | 3.26% | Genome-wide |
| AX-87879144 | ssa27 | 9354950 | 7.10E-07 | 6.15 | 2.13% | Genome-wide |
| AX-87990259 | ssa27 | 9602973 | 1.79E-05 | 4.75 | 1.60% | Chromosome-wide |
| AX-87107874 | ssa27 | 9875472 | 6.52E-12 | 11.19 | 4.05% | Genome-wide |
| AX-87716791 | ssa27 | 9885089 | 1.44E-10 | 9.84 | 3.54% | Genome-wide |
| AX-86960936 | ssa27 | 9972373 | 3.64E-08 | 7.44 | 2.62% | Genome-wide |
| AX-98308473 | ssa27 | 9974462 | 3.61E-05 | 4.44 | 1.49% | Chromosome-wide |
| AX-87784578 | ssa27 | 10016847 | 4.63E-06 | 5.33 | 1.82% | Chromosome-wide |
| AX-87784944 | ssa27 | 10025960 | 5.36E-07 | 6.27 | 2.18% | Genome-wide |
| AX-97869528 | ssa27 | 10049322 | 7.69E-06 | 5.11 | 1.74% | Chromosome-wide |
| AX-87388881 | ssa27 | 10176535 | 3.06E-07 | 6.51 | 2.27% | Genome-wide |
| AX-105314484 | ssa27 | 10177750 | 1.38E-06 | 5.86 | 2.02% | Chromosome-wide |
| AX-87889461 | ssa27 | 10192440 | 2.73E-06 | 5.56 | 1.91% | Chromosome-wide |
| AX-96309328 | ssa27 | 10348407 | 1.69E-14 | 13.77 | 5.03% | Genome-wide |
| AX-97902757 | ssa27 | 10393267 | 9.46E-15 | 14.02 | 5.13% | Genome-wide |
| AX-96395661 | ssa27 | 10512101 | 1.88E-06 | 5.73 | 1.97% | Chromosome-wide |
| AX-87311210 | ssa27 | 10654172 | 3.98E-06 | 5.40 | 1.85% | Chromosome-wide |
| AX-96197665 | ssa27 | 10749982 | 6.97E-06 | 5.16 | 1.76% | Chromosome-wide |
| AX-96156161 | ssa27 | 10750087 | 3.68E-06 | 5.43 | 1.86% | Chromosome-wide |
| AX-96395645 | ssa27 | 10781997 | 6.14E-06 | 5.21 | 1.78% | Chromosome-wide |
| AX-98308452 | ssa27 | 11167928 | 3.32E-07 | 6.48 | 2.26% | Genome-wide |
| AX-96309317 | ssa27 | 11168090 | 1.13E-07 | 6.95 | 2.44% | Genome-wide |
| AX-96197662 | ssa27 | 11185301 | 1.60E-10 | 9.79 | 3.52% | Genome-wide |
| AX-87474590 | ssa27 | 11191489 | 2.94E-06 | 5.53 | 1.90% | Chromosome-wide |
| AX-88058440 | ssa27 | 11681467 | 3.33E-10 | 9.48 | 3.40% | Genome-wide |
| AX-97886034 | ssa27 | 11723737 | 2.56E-05 | 4.59 | 1.54% | Chromosome-wide |
| AX-87205708 | ssa27 | 11973268 | 3.16E-07 | 6.50 | 2.27% | Genome-wide |
| AX-96420650 | ssa27 | 12208504 | 2.51E-05 | 4.60 | 1.54% | Chromosome-wide |
| AX-87259442 | ssa27 | 12499061 | 1.21E-05 | 4.92 | 1.67% | Chromosome-wide |
| AX-88275449 | ssa27 | 13298889 | 8.33E-06 | 5.08 | 1.73% | Chromosome-wide |
| AX-87941659 | ssa27 | 13563945 | 1.16E-06 | 5.94 | 2.05% | Chromosome-wide |
| AX-96156074 | ssa27 | 13570032 | 1.80E-06 | 5.75 | 1.98% | Chromosome-wide |
| AX-88293442 | ssa27 | 13627788 | 1.64E-07 | 6.78 | 2.37% | Genome-wide |
| AX-87649300 | ssa27 | 13896277 | 1.09E-07 | 6.96 | 2.44% | Genome-wide |
| AX-87232521 | ssa27 | 15079221 | 4.98E-06 | 5.30 | 1.81% | Chromosome-wide |
| AX-98317907 | ssa27 | 15160212 | 1.50E-06 | 5.82 | 2.01% | Chromosome-wide |
| AX-88045505 | ssa27 | 15161000 | 1.60E-05 | 4.80 | 1.62% | Chromosome-wide |
| AX-88266917 | ssa27 | 15161002 | 3.34E-05 | 4.48 | 1.50% | Chromosome-wide |
| AX-98317904 | ssa27 | 15161356 | 7.57E-06 | 5.12 | 1.74% | Chromosome-wide |
| AX-88093100 | ssa27 | 15622000 | 3.29E-09 | 8.48 | 3.02% | Genome-wide |
| AX-87670213 | ssa27 | 15923275 | 1.42E-05 | 4.85 | 1.64% | Chromosome-wide |
| AX-88138755 | ssa27 | 15986838 | 2.29E-05 | 4.64 | 1.56% | Chromosome-wide |
| AX-96310740 | ssa27 | 16179768 | 2.41E-05 | 4.62 | 1.55% | Chromosome-wide |
| AX-96231796 | ssa27 | 16507964 | 2.83E-09 | 8.55 | 3.05% | Genome-wide |
| AX-97879918 | ssa27 | 16528506 | 1.85E-05 | 4.73 | 1.59% | Chromosome-wide |
| AX-87140921 | ssa27 | 17265465 | 8.14E-07 | 6.09 | 2.11% | Genome-wide |
| AX-87877297 | ssa27 | 19804784 | 1.78E-05 | 4.75 | 1.60% | Chromosome-wide |
| AX-87336680 | ssa27 | 20250601 | 1.81E-06 | 5.74 | 1.98% | Chromosome-wide |
| AX-96121176 | ssa27 | 20445207 | 1.35E-05 | 4.87 | 1.65% | Chromosome-wide |
| AX-97902305 | ssa27 | 21423070 | 8.44E-07 | 6.07 | 2.10% | Genome-wide |
| AX-87990446 | ssa27 | 21428858 | 4.78E-06 | 5.32 | 1.82% | Chromosome-wide |
| AX-88140363 | ssa27 | 21497459 | 6.86E-07 | 6.16 | 2.14% | Genome-wide |
| AX-87633714 | ssa27 | 21528048 | 1.30E-05 | 4.89 | 1.65% | Chromosome-wide |
| AX-86925703 | ssa27 | 21645190 | 8.19E-06 | 5.09 | 1.73% | Chromosome-wide |
| AX-96451141 | ssa27 | 21646066 | 3.54E-06 | 5.45 | 1.87% | Chromosome-wide |
| AX-96438409 | ssa27 | 21649877 | 1.92E-08 | 7.72 | 2.73% | Genome-wide |
| AX-87708412 | ssa27 | 21914246 | 1.67E-05 | 4.78 | 1.61% | Chromosome-wide |
| AX-88309032 | ssa27 | 22175059 | 1.55E-06 | 5.81 | 2.00% | Chromosome-wide |
| AX-97880155 | ssa27 | 23148033 | 9.26E-06 | 5.03 | 1.71% | Chromosome-wide |
| AX-88244980 | ssa27 | 25362000 | 3.77E-07 | 6.42 | 2.24% | Genome-wide |
| AX-86978244 | ssa27 | 25448728 | 1.88E-07 | 6.73 | 2.35% | Genome-wide |
| AX-97877641 | ssa27 | 29299825 | 3.12E-05 | 4.51 | 1.51% | Chromosome-wide |
| AX-88003719 | ssa27 | 35744854 | 7.34E-06 | 5.13 | 1.75% | Chromosome-wide |
| AX-88277434 | ssa12 | 33472000 | 3.25E-05 | 4.49 | 1.50% | Chromosome-wide |
| AX-87027775 | ssa12 | 61391936 | 1.26E-05 | 4.90 | 1.66% | Chromosome-wide |
| AX-87253175 | ssa12 | 61443087 | 3.55E-05 | 4.45 | 1.49% | Chromosome-wide |
| AX-87704824 | ssa12 | 61694169 | 2.86E-05 | 4.54 | 1.52% | Chromosome-wide |
| AX-87796820 | ssa12 | 61701806 | 4.46E-06 | 5.35 | 1.83% | Chromosome-wide |
| AX-98299923 | ssa12 | 61702164 | 4.38E-06 | 5.36 | 1.83% | Chromosome-wide |
| AX-88191278 | ssa12 | 61702955 | 6.19E-06 | 5.21 | 1.78% | Chromosome-wide |
| AX-87018640 | ssa12 | 61703574 | 9.52E-06 | 5.02 | 1.70% | Chromosome-wide |

